# YAP independently regulates cell size and population growth dynamics via non-cell autonomous mediators

**DOI:** 10.1101/482836

**Authors:** Douaa Mugahid, Marian Kalocsay, Scott Gruver, Leonid Peshkin, Marc W. Kirschner

**Affiliations:** Department of Systems Biology, Harvard Medical School, Boston, 02115, MA, USA

## Abstract

The Hippo pathway, in which changes at the cell surface and in the extracellular environment control the activity of a downstream transcription factor, known as YAP in mammalian cells and Yorkie in Drosophila, has recently taken center-stage as perhaps the most important pathway in metazoans for controlling organ size. In intact tissues YAP activity is inhibited and the organ does not overgrow. When the organ is damaged, YAP is active and necessary for growth and regeneration to occur. The exact process by which YAP drives organ and tissue growth is not fully understood, although it is known to affect both cell size and cell number. Since cell size and proliferation are highly interdependent in many cultured cell studies, we investigated the role of YAP in the simultaneous regulation of both cell size and number. Our experiments reveal that YAP controls both cell size and cell proliferation by independent circuits, and that it affects each process non-cell autonomously via extracellular mediators. We identify that CYR61, a known secreted YAP target, is the major regulator of the non-cell autonomous increase in cell number, but does not affect cell size. The molecular identity of the non-cell autonomously acting mediator of cell size is yet to be identified.

## Introduction

It has long been appreciated that organ size is tightly regulated: it scales with body size, and in several cases organs can be regenerated after partial loss or injury (Penzo-Méndez and Stanger, 2015). An organ’s size is dictated by both the size and number of cells comprising it (Fankhauser, 1945). In adulthood, tissues maintain size largely through the proliferation and differentiation of tissue stem cells, and in some cases by the dedifferentiation of mature cells, followed by proliferation and re-differentiation. In some organs, the dedifferentiation-re-differentiation process can be skipped and the organs compensate for elevated demand by increasing cell size – such as skeletal and cardiac muscle, or by increasing both size (hypertrophy) and number (hyperplasia) – such as the β-cells of the pancreas (Anzi et al., 2018; Ernst et al., 2011). Such examples demonstrate that organ size is not simply set during development and passively maintained into adulthood, but is actively monitored to fit physiological demands. One organ that offers quite a dramatic example of size control is the liver. Although liver cells are largely quiescent in adults, the liver can regenerate completely after resection of up to 70% of its original mass (partial hepatectomy). This response is tri-phasic, and starts with an increase in the size of the remaining hepatocytes. Hypertrophy is followed by a period of cell proliferation, which produces daughter cells of normal size, and is terminated once the original mass is restored (Gilgenkrantz and Collin de l’Hortet, 2018). If liver cells are blocked from proliferation, liver size can still be restored after partial hepatectomy, simply by hypertrophy of the remaining cells (Diril et al., 2012). This indicates that, when the pathways inhibiting organ overgrowth are disrupted, pathways affecting both cell growth and cell proliferation can be activated to different degrees.

Based on genetic studies in Drosophila, the Hippo pathway was identified as having a particularly significant role in organ size regulation (Dong et al., 2007). It is now known to have similar functions in many systems including the mammalian liver, where it is plays roles in overgrowth inhibition (Dong et al., 2007) and regeneration after partial hepatectomy (Lu et al., 2018). The Hippo pathway is comprised of a cascade of intracellular kinases, activated in response to cell-cell contact. These kinases ultimately phosphorylate and inhibit the downstream transcriptional regulators YAP and TAZ, by causing their sequestration, and often degradation in the cytoplasm (Zhao et al., 2010). Upon tissue injury, the upstream kinases are inactive and YAP/TAZ translocates into the nucleus, where it mediates transcriptional changes that facilitate organ regrowth (Lee et al., 2014). Changes in the composition and physical properties of the extracellular environment can also lead to changes in YAP/TAZ activity (Dupont et al., 2011; Kim and Gumbiner, 2015). Several reports have shown that the activation of YAP increases cell proliferation *in vitro* and *in vivo* (Dong et al., 2007; Xin et al., 2011; Zanconato et al., 2015), while others have demonstrated that its effect is mainly on cell growth (Hansen et al., 2015). Each of these could provide a plausible mechanism for the role of the Hippo pathway in regulating organ size both by hyperplasia and hypertrophy.

In cultured cells, the regulation of cell size and number is strongly interdependent. While cell size is robustly maintained, it is under constant correction by regulation of cell cycle length and by compensatory changes in individual cells’ growth ((Ginzberg et al., 2018) and our unpublished results). Thus, a perturbation primarily affecting the cell growth machinery can affect cell size but at the same time have a clear effect on cell number. This intimate co-dependency of cell cycle and cell growth even in cultured cells may be obscuring our facile identification of many cell growth regulators as solely proliferation regulators. This seems particularly true of YAP, where there is already evidence that it can affect both cell size and number.

In this report, we have specifically looked at the underlying circuitry of YAP’s effects as a proliferation factor and as a regulator of cell size. We ask whether YAP affects both processes simultaneously and whether their regulation is inter-dependent. Our results suggest that YAP can affect both processes independently. Furthermore, we find evidence that both processes are driven by cell contact-independent, non-cell autonomous mediators. We further show that YAP regulates the transcription of a suite of extracellular proteins of various functional properties, including the CCN (CTGF, CYR61 and NOV) family protein, CYR61. CYR61 has no effect on cell size but mediates a significant part of the YAP-dependent increase in cell number in a non-cell autonomous manner. Changes in CYR61 levels can also explain the mechanosensitive YAP-dependent changes in cell number. We conclude that the non-cell autonomous effects on cell size are dependent on a separate secreted entity, which we have not yet identified.

## Results

### I) Overexpression of non-phosphorylatable YAP increases both cell number and size

To study the effect of YAP on cell size and number in culture, we generated HEK293 cells expressing nuclear mCherry (nmCherry) together with either nuclear GFP (nGFP), GFP-tagged wild type YAP (YAPWT) or a GFP-tagged phosphosite mutant YAP that is not phosphorylatable (YAP5SA). YAP5SA cannot be inhibited by the upstream kinases and as such is constitutively active. YAPWT is still subject to the same regulation as endogenous YAP. nmCherry allows facile counting of the number of cells in a culture at any given time. Throughout the manuscript all the YAP constructs to which we refer are GFP-tagged, but are referred to in an abbreviated form as: YAP5SA or YAPWT. In some experiments we use the area of the nucleus as a proxy for cell size as was previously demonstrated in (Ginzberg et al., 2018). With this experimental design, we can simultaneously track changes in cell number and average cell size over time (Figure S1).

The population growth rate in these cells decreased as cell density increased, and eventually plateaus (Figure 1A). This is a well-known phenomenon often referred to as density-dependent growth inhibition (Stoker and Rubin, 1967) or contact inhibition, and is captured by a logistic rather than exponential growth model. The exponential growth model approximates growth at low density but fails completely at high cell density (Figure S2A). Thus, all the growth curves presented are fits of our data to the solution of a logistic growth equation (Figure S2E), a differential equation first introduced by Verhulst (Pierre François Verhulst, 1845). At low population density, the model converges to exponential growth, but generally captures the bounded growth of populations at higher densities (Tsoularis and Wallace, 2002). In this model, changes in maximum cell number are described by *Ymax*, also known as the population carrying capacity; and the exponential growth rate is described by *k* (with units of reciprocal time), which reflects how fast a population of cells grows at low cell density. Our data shows that the population growth rate (k) and carrying capacity (Ymax) are independently regulated under different growth conditions, which we demonstrate in Figure S2 by varying the concentration of serum. Under such conditions, serum concentrations above 1 % changed k without changing Ymax. For that reason, we do not assume that the regulation of one affects the other and considered it necessary to discriminate between the factors that affect the two parameters. According to our data, the expression of YAP5SA increases both k and Ymax compared with the expression of nGFP alone (Figure 1B).

**Figure 1:**
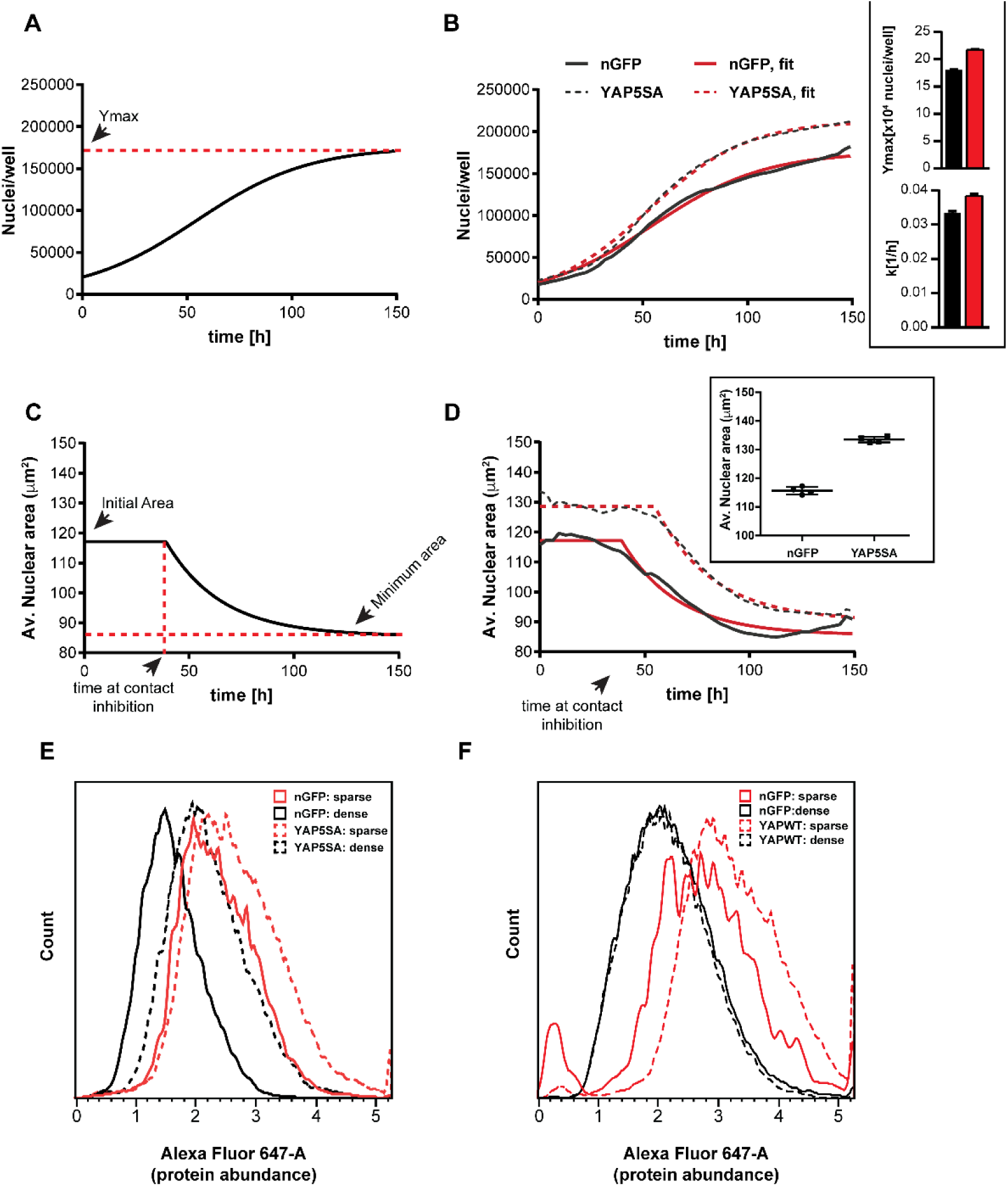
YAP5SA expression affects cell size and population growth dynamics. (A) Example of population growth dynamics in HEK293 cells in cultures seeded at low density. Ymax is the carrying capacity of the population according to the logistic growth model. (B) Constitutively active YAP (YAP5SA) increases the rate of growth (k) and carrying capacity (Ymax) of cells vs. nuclear GFP (nGFP) controls. (C) Nuclear area of HEK293 cells is initially constant, but exponentially decays to a minimum as cell number increases. (D) Nuclear area is larger in YAP5SA cells vs. controls, but still decreases exponentially as cell density increases. (E) Protein content is higher in YAP5SA cells vs. controls at high and low density. (F) Protein content is higher in cells overexpressing wildtype YAP (YAPWT) vs. nGFP controls at low density but not at high density. A, C: In black: the average of 4 replicates; in red: the fit to a logistic model. B, D: n=4; mean±SD.

With regard to cell size, low density cultures (< ~10,000 cells/well in a 96-well plate) initially maintained a constant average nuclear area up to a critical density, after which the nuclear area decreased to a minimum following a roughly exponential curve (Figure 1C). YAP5SA cells displayed an initial average nuclear area that was significantly larger than nGFP controls (~17%). A change of such magnitude would translate to ~30 % more volume (assuming cell to be spherical). This indicates that YAP increases cell size at low cell density as has been previous reported (Hansen et al., 2015). However, as the number of cells continued to grow, the nuclear area of YAP5SA cells also decreased to a minimum where the difference in size compared with controls is much smaller (Figure 1D). Since size at the later time points seems to be a complex product of initial size and the achieved Ymax, we focused on changes in nuclear area (cell size) in low density cultures.

To validate that the changes in nuclear area reflected changes in cell size, we measured protein mass directly in individual cells using a covalent dye specific for lysine groups on the protein. Since protein mass represents the most significant increase in dry mass during growth (Winick, 1968), we consider changes in protein content a good proxy for changes in size. Though Ginzberg *et al*. (Ginzberg et al., 2018) demonstrated a good correlation between protein content and nuclear area in their cell type, there is no a priori reason why they should be related in all cells.

We found that average protein content per cell was ~30 % higher in populations expressing YAP5SA (Figure 1E) than in nGFP controls, supporting the estimates made based on the change in nuclear area (Figure 1D). Since YAP5SA increased cellular protein content throughout all the phases of the cell cycle (Figure S3B), we ruled out that the observed changes were merely due to an increase in the fraction of cells in S or G2/M (which are on average larger than cells in G1; Figure S3A). This analysis was done by co-staining for total protein and DNA content and comparing the distribution of protein content in cells based on their position in the cell cycle. We also demonstrated that cellular protein content was reduced in high density cultures (Figure 1E). This reinforced the conclusions drawn from the changes in nuclear area showing that YAP5SA cells, like nGFP controls, are still subject to density-dependent changes in cell size (Figure 1D). Similar experiments using YAPWT-expressing cells demonstrated that YAPWT overexpression only increased cell size in low density cultures, but not at high density (Figure 1F). This is expected because at high density YAPWT should be inactivated by the upstream kinases of the Hippo pathway, while YAP5SA should not. One puzzling observation was that cells expressing nGFP were themselves larger than mock-transfected controls which we further discuss below.

### II) YAP mediates non-autonomous changes in cell size and number

We wished to address the puzzling size increase in nGFP-expressing cells, which we thought might be due to some subtle uncorrected differences in the culture conditions. To control for effects of cell density and possible heterogeneity in the culture conditions, we compared the size of cells expressing GFP-tagged YAP5SA or nGFP with WT cells co-cultured in the same well (Figure 2A). This set-up allowed us to use GFP to differentiate between the two populations, even though they are intermixed. Surprisingly, we found that in co-culture the differences in protein content between nGFP-expressing and WT cells were maintained (Figure 2B, solid lines). The observed increase in size due to GFP expression (~20%; Figure 2B; solid line, red) is smaller than the effect of YAP5SA but reproducible (Figure 2B; dashed line, red). The increase in protein content cannot be accounted for by the mass of GFP expressed by these cells (~0.2 % of total protein; data not shown), suggesting that GFP overexpression is not functionally inert. Nevertheless, we observed no difference in the size of GFP-negative cells (WT) and those expressing GFP-tagged YAP5SA in co-culture.

**Figure 2:**
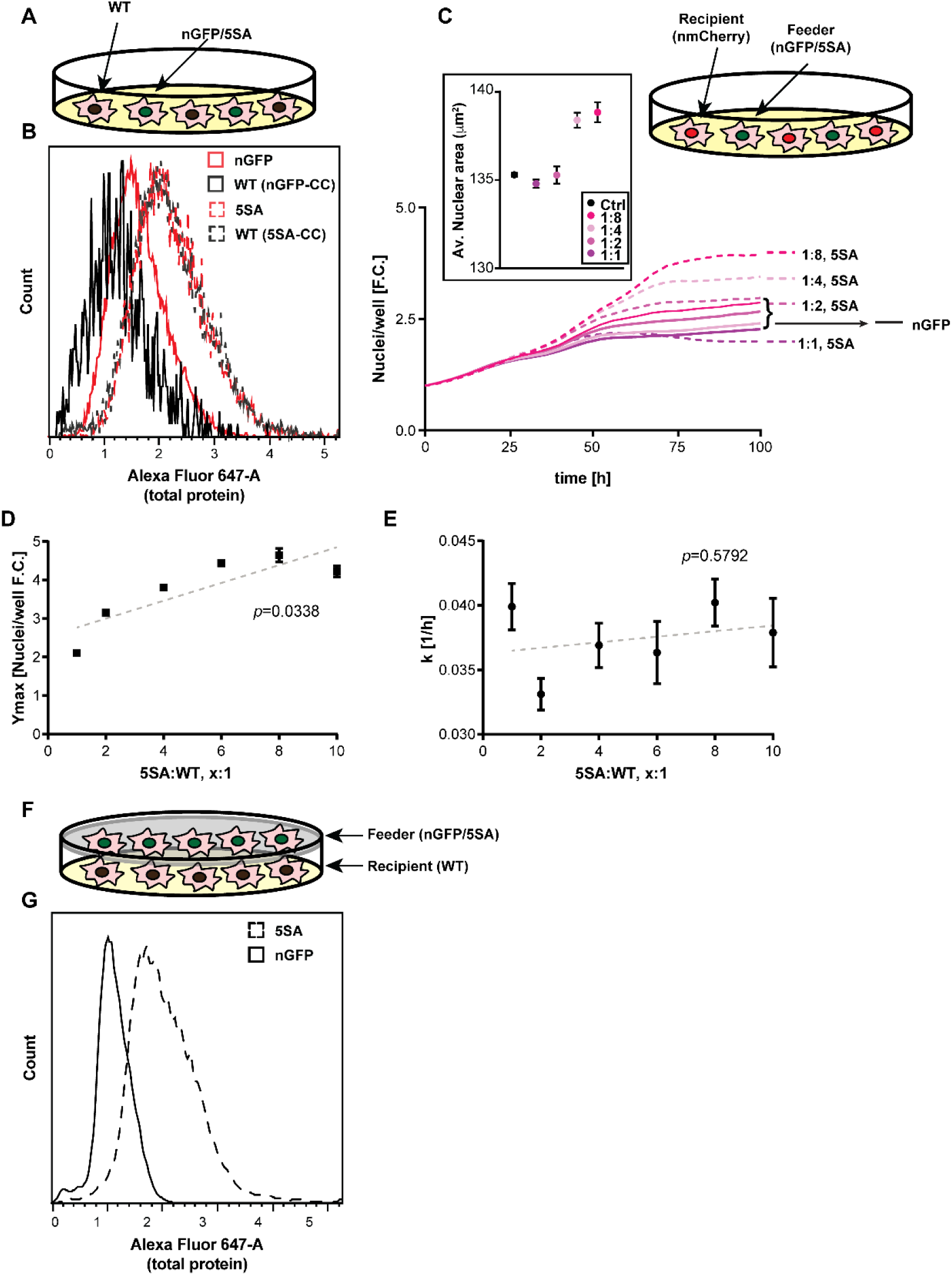
YAP5SA increases cell size and the carrying capacity of cells cell non-autonomously. (A) Co-cultures of nGFP or GFP-tagged YAP5SA intermixed with WT cells. (B) nGFP-expressing cells (solid red line) co-cultured with WTs (solid black line) have higher average protein content. Both GFP+ (red dotted line) and GFP-cells (black dotted line) in YAP5SA-co-cultured cells are larger than cells in the nGFP co-cultures. (C-E) Increasing the fraction of YAP5SA cells in co-cultures with nuclear mCherry cells (nmCherry) increases the carrying capacity (Ymax) of the nmCherry cells as well as their nuclear area, but does not affect population growth rates (k); *p-values* indicate the likelihood that the slope is non-zero. (F) WT cells co-cultured with YAP5SA or nGFP cells on a Transwell™ membrane exchange medium components without physical contact. (G) WT cells co-cultured with YAP5SA cells have higher protein content than those co-cultured with nGFP cells. n=5; C: mean±SD (C). D, E: mean±SEM.

Most surprising is, when the nGFP and YAP5SA co-cultures were examined at comparable densities, cells expressing nGFP and YAP5SA, as well as those co-cultured with YAP5SA but not expressing any transgene, showed an increase in size over the WT cells co-cultured with nGFP cells (Figure 2B). While, the smaller effect of nGFP expression was limited to the cells expressing nGFP and hence appeared to be cell autonomous (Figure 2B; solid lines; red vs black), the increase mediated by YAP5SA is clearly non-cell autonomous and is also much larger in magnitude (Figure 2B; dashed lines). These results demonstrate that even though the expression of YAP5SA is obviously a cell autonomous process, the effect of the transfected YAP5SA is shared equally with the untransfected cells in the same well.

To investigate whether YAP expression affected not only cell size but also cell number in a non-autonomous manner, we co-cultured cells expressing nmCherry “apparent recipients” with cells expressing either YAP5SA or nGFP alone “apparent feeders” (Figure 2C). The two cell types were well-mixed while varying the ratio between the recipient and feeder cells. We found that nmCherry cells co-cultured with YAP5SA cells had a larger nuclear area and their population grew to a higher number when YAP5SA cells were in large excess (four to eight times as many), compared with those co-cultured with nGFP controls (Figure 2C, D). The population growth rate at low density (k) did not significantly change with the number of feeder cells (Figure 2E). Ymax, however, increased ~ 300 % at a 1:4 ratio of nmCherry:YAP5SA cells (Figure 2D) and even more so at a ratio of 1:8. The effects on cell size were smaller, with changes in nuclear area reflecting an increase corresponding to ~ 5 % in size (Figure 2C). Attempts to drive nuclear area by increasing the fraction of YAP5SA cells to more than four fifths of the population were unsuccessful. Note that cells in these experiments were seeded at relatively high densities, which we believe may explain the smaller changes in size under these conditions. This suggests that the factors affecting individual cell growth over the length of the cell cycle (size) are different from those affecting the population’s capacity for growth at high density (Ymax), or at least suggests that the sensitivity of these processes to the same regulator is different.

Non-cell autonomous effects could be due to the physical interaction of cells, such as cell-cell contact, or could be mediated by the diffusion of molecules (including their depletion from culture medium). However, we reasoned the general depletion of a component from the medium is less likely to be the explanation, since that would imply that cells would grow larger over time as the factor continued to decrease. However, our measurements indicate this is not the case (Figure 1D, E). To investigate whether these effects required cell-cell contact, HEK293 cells were seeded on the bottom of a well, physically separated from a population of feeder cells expressing YAP5SA or nGFP in a Transwell^®^ chamber. The feeders were seeded on an insert that allowed the free diffusion of small molecules and peptides but not cells (Figure 2F). This setup allowed both cell types to share and exchange medium components without allowing their physical contact. Cell size, measured by protein content, was then compared between the cells co-cultured with each feeder cell type. Cells co-cultured with YAP5SA cells had higher levels of protein on average compared with those co-cultured with nGFP controls (Figure 2G), suggesting that YAP’s non-cell autonomous effect on cell size is likely mediated by a soluble factor produced by YAP5SA-expressing cells. This spurred us to look for specific factors secreted by the YAP5SA cells that could stimulate growth at the cellular and population level.

### III) A large fraction of proteins affected by YAP5SA overexpression are extracellular or membrane proteins

The simplest interpretation of these results is that these non-cell autonomous effects are mediated by secreted proteins that are transcriptional targets of YAP. We therefore tried to identify these factors using RNA-Seq and mass spectrometry (Supplementary table S5). Our analysis revealed that extracellular proteins were a surprisingly large portion of the those differentially expressed in response to YAP5SA (Figure 3B). In some cases, large changes in the protein level were not accompanied by correspondingly large changes in the mRNA level (Figure 3A). Among the most prominently changed extracellular proteins were CYR61 and CTGF, which were elevated at both the protein and RNA level. Both proteins are members of the CCN family of matricellular proteins, non-structural proteins that reside in the extracellular matrix and regulate cellular function, often by binding to and affecting the potency of other growth factors (Holbourn et al., 2008). Both genes have been previously used as robust readouts at the transcriptional level for YAP activity (Dupont et al., 2011; Yimlamai et al., 2014). Nevertheless, their contribution to the YAP-dependent changes in cell behavior has never been clarified. Therefore, we decided to probe deeper to evaluate whether either – or both – of these CCN proteins mediate YAP’s effects on cell size and/or population growth. Amphiregulin, a secreted signal related to EGF, had previously been identified as a potential mediator of YAP-dependent cell non-autonomous growth in 3D cultures of MCF10 cells (Zhang et al., 2009). In our experiments, Amphiregulin was expressed at a very low level (FPKM <1 compared with FPKM >100 for CYR61; neither could we identify Amphiregulin by mass spectrometry). This suggested that factors other than Amphiregulin are more likely involved in the cell size and population growth changes we observed.

**Figure 3:**
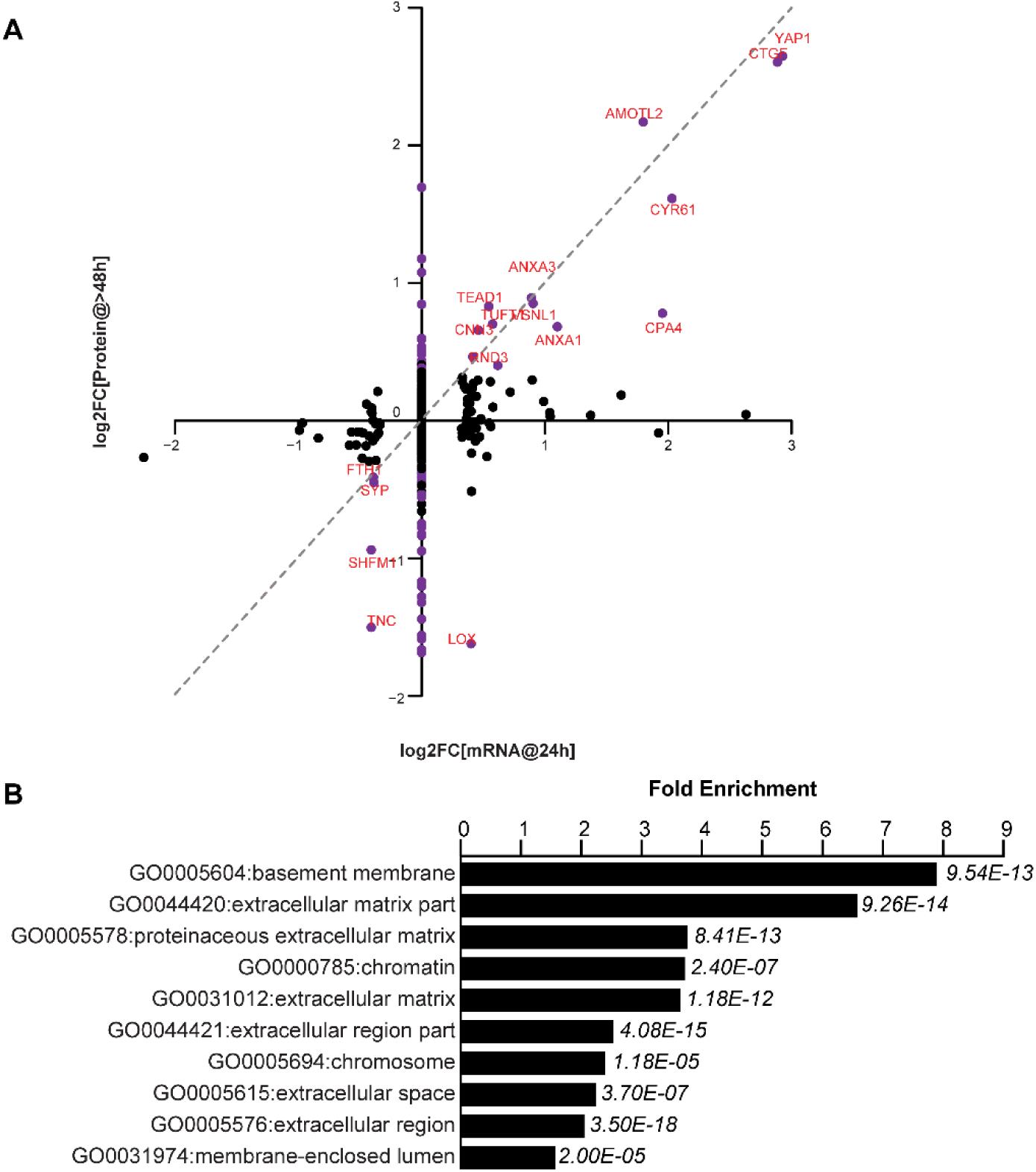
YAP-dependent changes in protein and mRNA expression levels. (A) Scatter plot of changes in protein (y-axis) vs. mRNA levels (x-axis) upon YAP expression. Dots in purple indicate proteins have a log2[fold change] (log2FC) more than 2 standard deviations away from the mean. Dotted line indicates a 1:1 change in protein vs. mRNA levels. (B) The 10 most significantly enriched cellular compartments in which regulated proteins reside (according to fold enrichment relative to all identified proteins). Benjamini-Hochberg *adjusted p-values* are indicated in italics.

### IV) YAP target, CYR61, causes an increase in the number of cells in culture but has no measurable effect on cell size

To determine whether CYR61 and CTGF affect non-cell autonomous changes in cell size and/or population growth, we used neutralizing antibodies to inhibit their extracellular activity in co-cultures seeded at a ratio of 1 WT:6 YAP5SA cells. Neutralizing CTGF had a small effect on cell proliferation, mildly decreasing k (~12 %), and mildly increasing Ymax (~12 %) of nmCherry cells (Figure 4 A–C). On the other hand, neutralizing CYR61 significantly decreased Ymax (~50 %) but increased k (~200%) in a dose-responsive manner (Figure 4 D–F). These findings support a role for CYR61 in the non-cell autonomous, YAP-mediated increase in Ymax. Although these results do not reflect on the potency of these molecules, they attest to the relative importance of CYR61 compared with CTGF in our cultures. Our RNA-Seq data suggest that the amount of CYR61 (FPKM > 100) produced in these cultures is considerably greater than CTGF (FPKM < 50). Consistent with the difference in expression levels, increasing the amount of antibody had an enhanced effect in the case of anti-CYR61 but not anti-CTGF. Neither neutralizing CTGF nor CYR61 affected nuclear area, suggesting that some other factor(s) are responsible for the non-autonomous effects of YAP on cell size (Figure 4 G, H).

**Figure 4:**
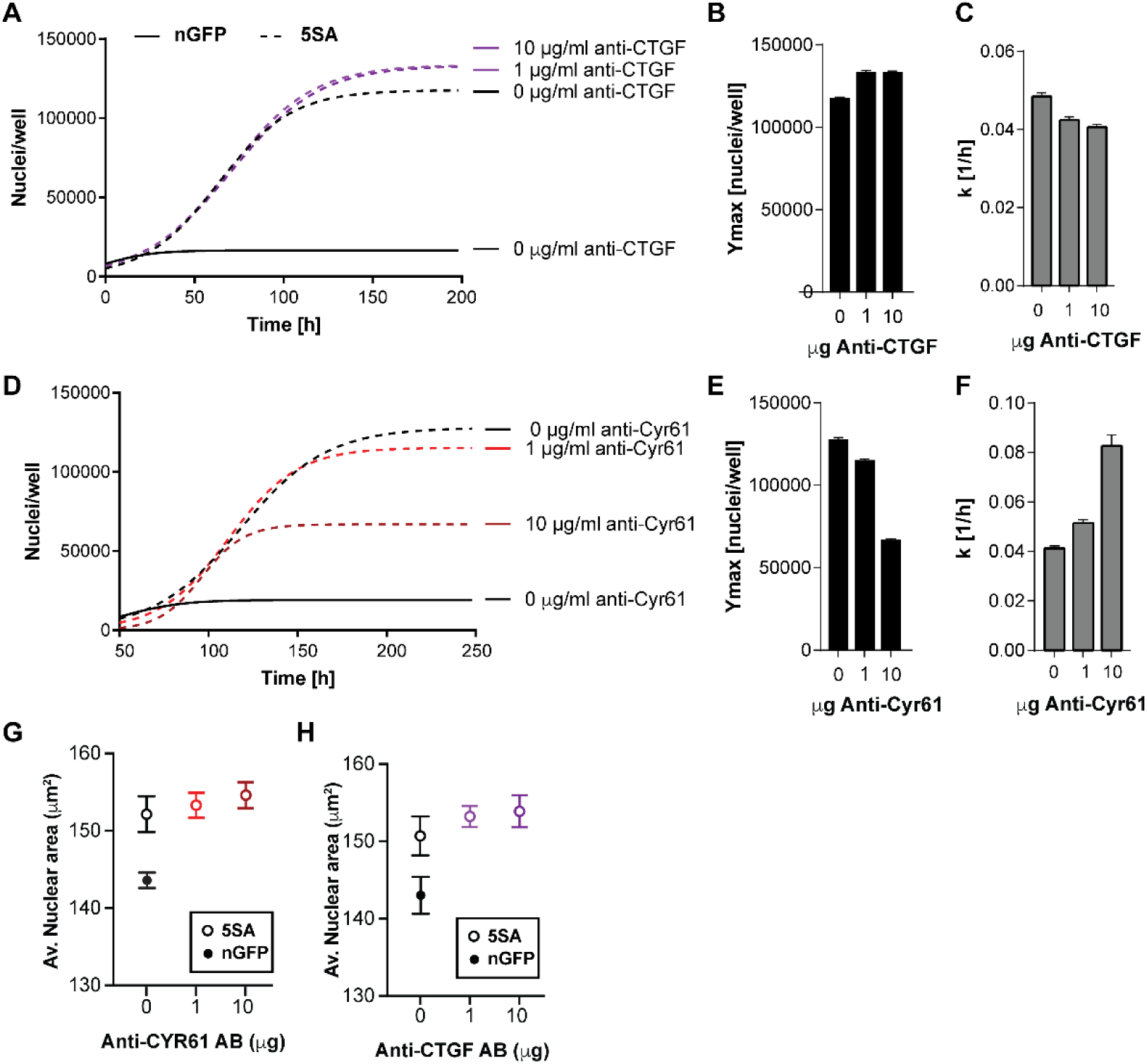
The effects of CTGF and CYR61 neutralization on the behavior of WT cells co-cultured with YAP-5SA cells at 1:6. (A-C) Neutralizing CTGF in co-cultures of nuclear mCherry (nmCherry) increases the nmCherry cells’ carrying capacity (Ymax), while decreasing their rate of growth (k). (D-E) Neutralizing CYR61 in these cultures has opposing effects of larger magnitude vs. CTGF. (G, H) Neutralizing antibodies have no effect on cell size. B, C, E, F: n=5; mean±SEM. G, H: n=5; mean±SD.

When treated with exogenous recombinant CYR61, both cells expressing YAPsh (YAPKD) and empty vector controls responded by increasing Ymax without a change in nuclear area (Figure 5). This confirms that CYR61’s non-cell autonomous influence on population growth – specifically the carrying capacity (Ymax) – does not require active YAP in the responding cells. Furthermore, CYR61 depletion or addition had no measurable effect on cell size.

**Figure 5:**
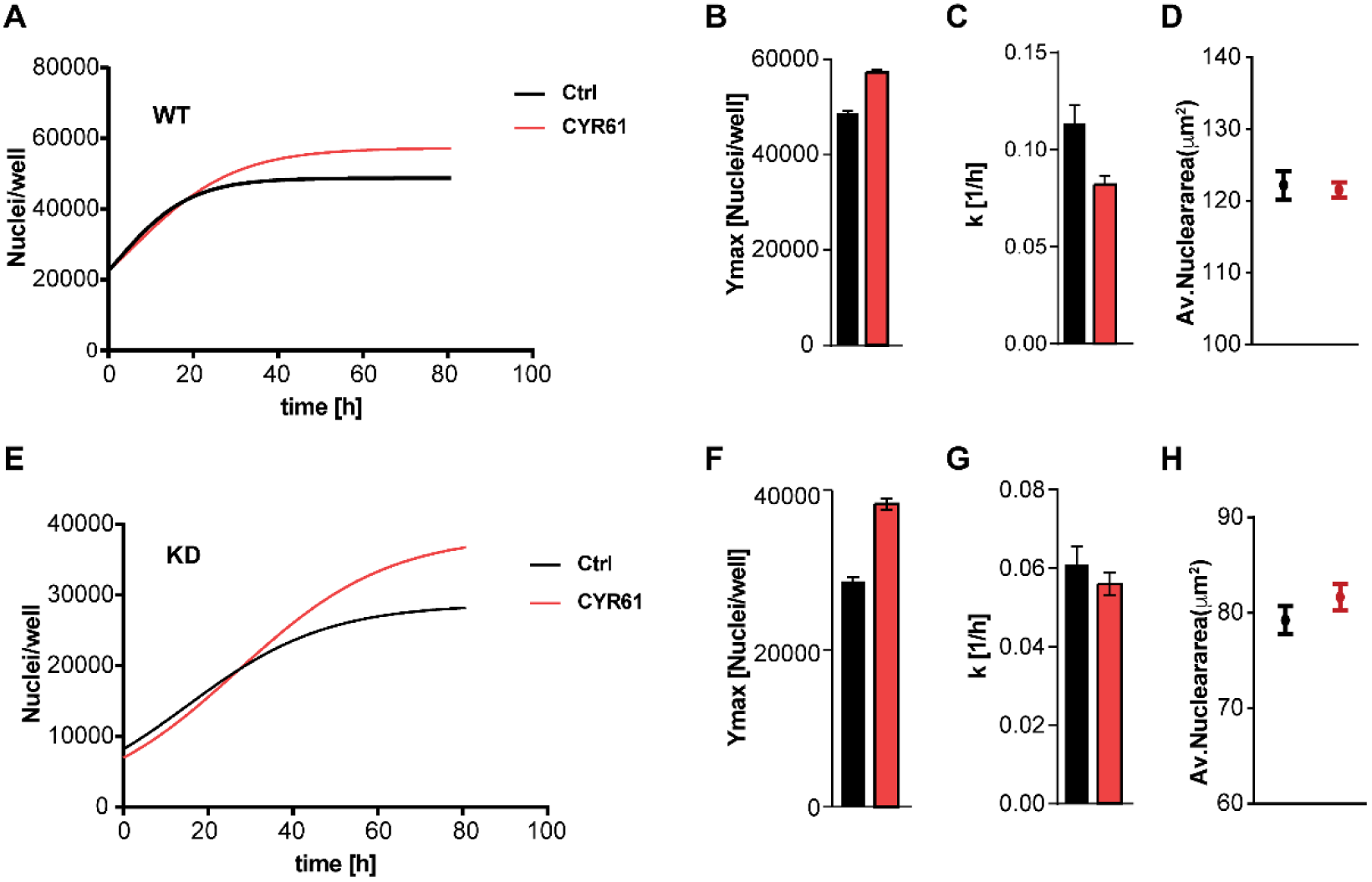
Exogenous CYR61 affects cell growth irrespective of YAP expression levels. (A-D) Treating WT cells with CYR61 increases their carrying capacity (Ymax) and decreases their growth rate (k) without altering initial nuclear area. (E-H) YAPKD cells respond to exogenous CYR61 by increasing Ymax, decreasing k with little effect on nuclear area. B, C, F, G: n=5; mean±SEM. D, H: n=5; mean±SD).

### V) Do the mechanical properties of the cell substrate convey some of the cell non-autonomous effects of YAP?

In a number of studies the stiffness of the substratum has been shown to act through YAP to mediate changes in cell behavior (Dupont et al., 2011; Halder et al., 2012). Thus far, we have focused our studies on the role of secreted molecules downstream of YAP in mediating non-cell autonomous changes in cell number and size. We recapitulated some of these effects most convincingly by mixing varying fractions of YAP5SA cells with WT cells. Since increased stiffness also results in an increase in the fraction of cells expressing nuclear YAP in a population (Dupont et al., 2011; Elosegui-Artola et al., 2017), we examined the role of substrate stiffness in controlling growth kinetics and cell size. We first cultured YAP5SA cells on Collagen I-coated substrates of increasing stiffness (8-50 kPa). We found no change in nuclear area or population growth dynamics (Figure 6A-D). However, in these homogeneous cultures all cells in each condition overexpressed active YAP, and YAP-dependent transcription was likely the same regardless of stiffness. This means that, in this setup, cells across all conditions might have been producing saturating levels of CYR61 and the factor(s) controlling cell size. To retreat from this potential state of saturation, we repeated the experiment with controls expressing empty vector and knock down (YAPKD) cells expressing 20 % the amount of YAP in these controls (Figure 6E). Consistent with our previous observations, the average nuclear area of YAPKD cells was smaller than controls. While stiffness did not change initial nuclear area in either cell line under these conditions (Figure 6F), it affected population growth (Figure 6G). Ymax increased as a function of stiffness in both YAPKD and control cell lines, although YAPKDs had a lower carrying capacity than controls on comparable substrates (Figure 6 H). On the other hand, growth rate, k, was slightly higher in YAPKD cells compared with controls on substrates of elastic modulus ≥ 25 kPa (in fact slightly higher), but was invariant with stiffness in the lower ranges in the YAPKDs. The observed increase in Ymax and decrease in k on different substrates can be simply explained by an increase in the fraction of cells with nuclear YAP. The picture is simpler to explain if we think of the effects to be mediated by a secreted factor like CYR61. The stiffer the substrate, the more cells expressing nuclear YAP, the more CYR61 is produced. This stiffness-dependent increase in CYR61 could explain both the increase in Ymax and the decrease in k (compare Figures 6G-1 to Figures 4D-F and 5). Although the fraction of cells expressing nuclear YAP is the same in both controls and YAPKDs, the level of CYR61 produced by the YAPKD cells is less, thus the lower Ymax and higher k. This would also explain why YAPKD cells are not responsive to stiffness in the ranges below 25 kPa. We confirm that CYR61 transcription is significantly higher in cells cultured on stiffer substrates after reanalyzing stiffness-dependent expression data from GSE102350 (Chang et al., 2017) (Figure 6J; Supplementary Table S6). Both Cyr61 and CTGF (Figure 6J, K) are among the transcripts most significantly regulated in response to stiffness, as would be expected in response to increased YAP activity.

**Figure 6:**
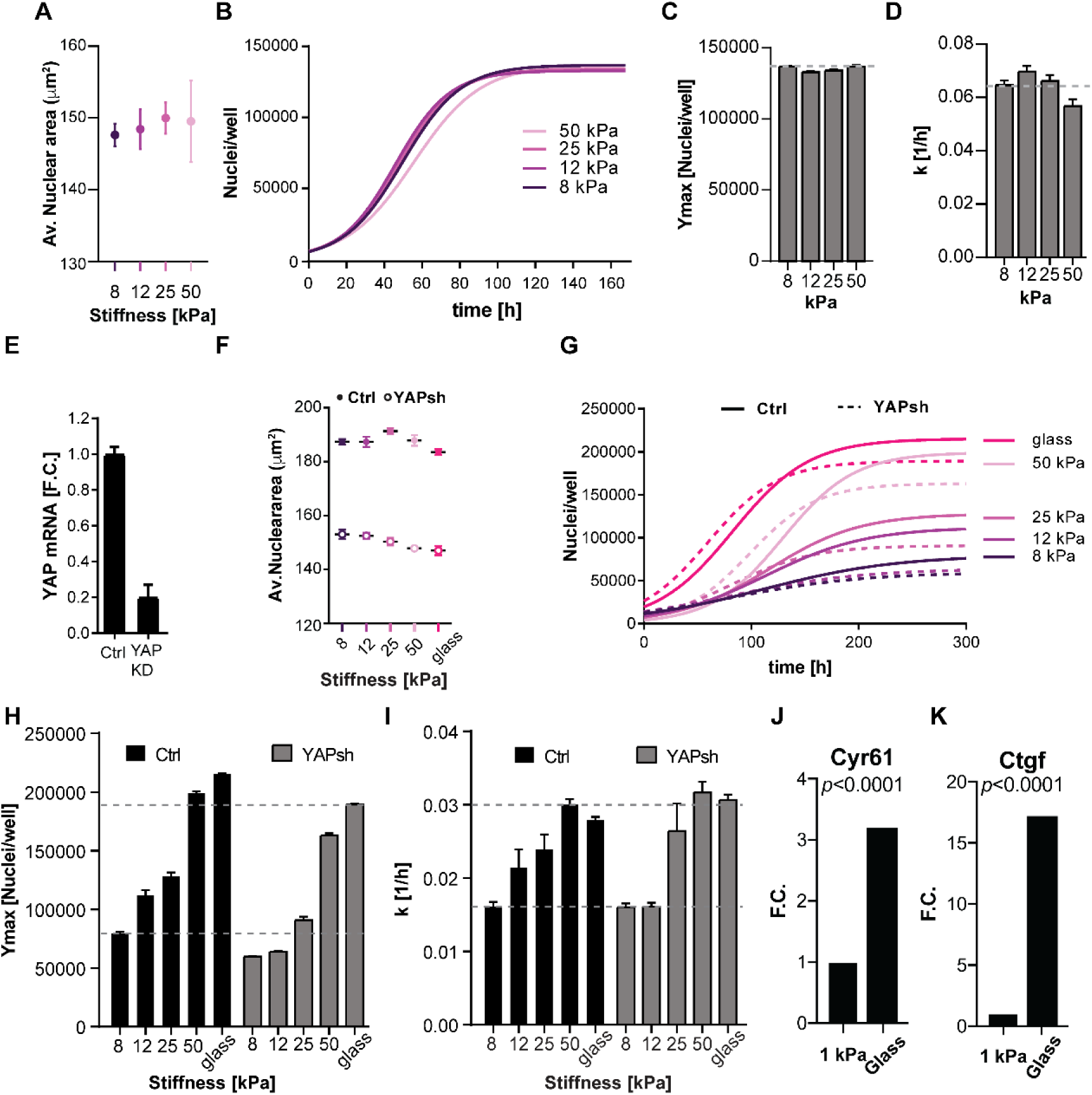
YAP affects substrate stiffness-dependent changes in population growth. (A-D) YAP5SA-expressing cells grown on substrates of increasing stiffness display no stiffness-dependent changes in initial nuclear area or population growth. (E) HEK293 cells expressing anti-YAP shRNA (YAPKD) express 80% less YAP mRNA vs. controls (ctrl). (F) The nuclear area of YAPKDs is smaller than ctrls, but stiffness does not significantly affect nuclear area in either cell line. (G) Fitted growth curves of control and YAPKD cells grown on Collagen I-coated substrates of increasing stiffness (8 kPa to glass). (H) The carrying capacity (Ymax) of WT and YAPKD cells increase as substrate stiffness increases. YAPKDs have a lower carrying capacity than controls grown on similar substrates. (I) The rate of growth (k) of WT and YAPKD cells increases with substrate stiffness. In YAPKDs, k is not affected on soft substrates (< 25 kPa) but is higher than controls on substrates of stuffniess 25 kPa and above. (J, K) Fold change (F.C.) in mRNA levels of Ctgf and Cyr61 levels in cells cultured on glass vs a 1kPa substrate (data from GEO dataset GSE102350). A-D: n=5; E-I: n=4. A, E, F: mean±SD; C, D, H, I: mean±SEM.

## Discussion

In the early days of physical chemistry, the laws of solutions could only be formulated satisfactorily at low concentration. Similarly, in early studies of cell growth the focus was on unicellular organisms or cells grown at low density. In both cases, high density meant having to deal with complex interactions of the individual components, the space they occupy, and effects on the solvent or the medium. At low concentration cell proliferation could be described by a simple differential equation, exponential growth, while at high concentrations there was no such simple model. Instead at high density cell growth slows, eventually stops and then declines. Interest in growth at high cell density was piqued when biologists made 2D cultures of animal cells on petri dishes. For example, chicken embryo fibroblasts grown in such a manner generate a monolayer and stop growing (Robert A. Weinberg, 2007). It is unlikely that this was simply depletion of the medium, for when Rous Sarcoma Virus was added the quiescent cells were transformed and proliferated, overgrowing the monolayer and producing three dimensional “foci” on top of the monolayer (Groupe and Manaker, 1956). There are two phenomena here to consider, the still unexplained tendency of many cells to bump into each other and stop growing and the tendency of a few infected or transformed cells to overcome whatever inhibitions were observed in the uninfected monolayers. With the passage of 50 years these phenomena have neither been completely resolved nor completely forgotten. The problem of cell interactions is still a serious topic in cancer (Kamińska et al., 2015) and more recently has emerged in many other biological settings, such as the cell behavior in regeneration or in stem cell niches (Lane et al., 2014).

More recently, interest in the reciprocal interactions of proliferating and resident cell populations was stirred by the discovery of the Hippo pathway, originally discovered in Drosophila. The major effector of the pathway is the transcription regulator YAP, which is formally considered an oncogene, because it stimulates growth (Dong et al., 2007). Over-activating YAP results in a breakdown of some of the usual barriers to growth in complex tissues and in high density cell cultures (Zhao et al., 2007). An interesting feature of the pathway is its sensitivity to alterations in the nature of the substratum, such as its stiffness (Dupont et al., 2011; Kim and Gumbiner, 2015). It is still generally assumed that the aggressive behavior of the cells is governed in a cell autonomous manner – like the original Rous transformation mentioned above. Another, and more unique feature of YAP is its control of organ and cell size. Cell growth and proliferation are two widely-studied phenomena common to all of life, but their coordination, which ultimately governs size control, is not well understood. We know that drugs or mutants that inhibit growth usually evoke a compensatory delay in cell division, thus preserving size. Similarly, perturbations that slow down the cell division cycle are generally compensated by a slowing down of growth that also preserves size (Ginzberg et al., 2018). YAP’s ability to perturb this well buffered balance of growth and division of cells is fascinating and largely unexplored.

We approached the role of YAP in size and proliferation control by formally considering the population’s behavior across the full growth range – from the early exponential phase to the plateau. We found that the growth rate (k) of cells expressing the activated YAP gene, YAP5SA, was slightly elevated (~14 %) compare to wild type. But the maximum carrying capacity of the population was more greatly affected, increasing about 35 %. The size of cells, reflected in changes in nuclear area and average protein content, was increased by as much as 40 % in some instances. Although these characteristics are not usually measured, they seem consistent with what might be expected for a dominant oncogene, albeit one that overrides size control. The previous reports of the effects of YAP on cyclin E and cyclin D supported this assumption (Dong et al., 2007; Shu and Deng, 2017; Tapon et al., 2002).

Although it was known that YAP increased the levels of several secreted gene products, it was not known that its major effects on cell size and number would be non-cell autonomous. In co-cultures of YAP5SA-expressing cells and wild type cells (WT), both cell types increased *equally* in size, indicating that the entire effect – within our error of measurement – was non-cell autonomous. In addition, the number of cells at the growth plateau, the carrying capacity of the population (Ymax), increased in WT cells in proportion to the number of YAP5SA cells in the culture, again indicating the regulation of Ymax was also non-cell autonomous. Only the initial exponential rate of growth in the WT cells, k, was unaffected by the co-cultured YAP5SA cells. Though increasing the fraction of YAP5SA cells in co-cultures increased both cell size and number, the effect on size saturated at lower levels. These findings argue that the striking effects of expression of activated YAP on the carrying capacity of the population and the size of the cells, were mediated by diffusible signals. Presumably, cells with active YAP are responding to autocrine rather than cell autonomous signals, while neighboring WT cells are responding to secreted (paracrine) signals.

Transcriptomic and proteomic analysis identified mostly membrane and extracellular proteins as those affected by the overexpression of YAP5SA. One of the strongest proteins to be differentially expressed was CYR61, a member of the CCN group of matricellular proteins, known to be involved in tissue remodeling (Chen et al., 2001), inflammation and embryogenesis (Katsube et al., 2009; Latinkic et al., 2003). In our experiments, recombinant CYR61could recapitulate the increase in Ymax measured by our co-cultivation experiments, but not the effect on cell size. Moreover, we show that substrate stiffness’ effects on population growth are YAP-dependent and can be explained via changes in CYR61 production. Stiffness did not significantly alter cell size irrespective of YAP levels. We thus conclude that cell non-autonomous signals mediated by YAP are responsible for both population growth and cell size effects but that the secreted factors causing them are different. It is worth noting that CTGF mRNA and protein levels are also increased in response to YAP5SA expression. Though CTGF and CYR61 are members of the CCN family of matricellular proteins, our data suggest that they have opposing effects on population growth but that the levels of CTGF in our cultures are small compared with CYR61. We believe this difference explains why the observed effects on population growth are overwhelmingly those of CYR61. We did not find evidence that the classic size regulator, such as IGF1/2, are produced to any appreciable degree by our cells. The identity of the potential secreted *size* mediator remains elusive.

CCN proteins themselves are not known to bind to any receptor tyrosine kinases, but rather bind to other growth factors thereby affecting the availability and/or affinity of growth factors for their receptors (Leask and Abraham, 2006). It is thus not surprising that cultivating cells at serum concentrations of less than 1 % decreased Ymax in our cultures (Figure S2), or that YAP5SA cells cultured under similar conditions proliferated at rates indistinguishable from controls (Figure S4A). We therefore surmise that an additional factor must be present in the serum at an adequate concentration for CYR61 to function. What this factor is, however, is another open question. Since we find that size is also affected by the level of serum (Figure S4 B), it is possible that the YAP-dependent size regulator is another metricellular protein that acts in concert with some growth factor in the serum.

Work by Tian and colleagues (Tian et al., 2013) had previously demonstrated that a high molecular weight hyaluronic acid produced by naked mole rate fibroblasts contributes to early contact inhibition; the equivalent of a decrease in the population’s carrying capacity. This form of hyaluronic acid acts via the upstream Hippo component NF2, which inhibits YAP activity. Combining this conclusion with our results suggests the importance of extracellular cues as part of the regulatory program for cell and organ size. YAP seems to play a role in sensing these extracellular changes and, in turn, signaling to adjacent cells to proliferate and grow and – ultimately – to further remodel the extracellular matrix. Given the non-cell autonomous nature of YAP signaling, local changes can affect widespread cellular processes such as regeneration and proliferation on a scale much greater than that accessible to the autonomous effects of single cells. Such a mechanism could explain how, *in vivo*, the activation of YAP in a small fraction of cells at the site of injury could engage a wider portion of the tissue or even a whole organ to participate in tissue repair; or even recruit the participation of other cell types. In considering the effects of secreted signals driven by activated YAP on cell number and size, we are probably only skimming the surface of roles that YAP may play in more complex physiological responses. It is significant that in the accompanying paper by Hartman and colleagues (doi: https://doi.org/10.1101/481127), the authors show that YAP mediates the reprogrammability to iPS cells both autonomously and non-autonomously. They demonstrate that the efficiency of the process is also non-cell autonomously dependent on matricellular proteins, particularly CYR61, transcriptionally regulated by YAP.

## Supporting information

## Acknowledgements

We thank Mike Gage for his meticulous assistance with the preparation of the protein samples for mass spectrometry and his overall management of project resources. We thank Jodene Moore of the SysBio FACS facility for her indispensable help setting up FACS analyses on the facility machines. We also thank the BPF Next-Gen Sequencing Core Facility at Harvard Medical School for their expertise and instrument availability that supported this work. Finally, we are grateful to Shangqin Guo and Amaleah Hartman for the stimulating discussions and thoughtful critique of this paper.

## Author contributions

MWK and DM wrote the manuscript and designed the experiments. DM, MK and SG performed the experiments. DM, MK and LP performed the data analysis. All authors have reviewed and approved the contents of this manuscript.

## Materials and methods

### Tissue culture

PEGFP-C3-YAP5SA plasmids were generated from the pEGFP-C3-hYAP1 plasmid gifted from Marius Sudol (Addgene plasmid # 17843 (Basu et al., 2003)). YAP amino acids S61, S109, S127, S164, S381 were all mutated to Alanine using the QuikChange II XL Site-Directed Mutagenesis Kit (Agilent). The following nuclear localization signal atggatccgaagaaaaaacgtaaaggccgtatggatccgaagaaaaaacgtaaaggccgt was appended to the 3’-end of GFP in the pAcGFP1-C3 vector (Clontech) to ensure its nuclear localization. Dox-inducible HEK293 cell lines of nGFP or GFP-tagged YAP5SA were generated by sub-cloning either construct into the pcDNA™5/FRT expression vector (Invitrogen) then co-transfecting the generated plasmids with the pOG44 Flp-Recombinase Expression Vector to ensure recombination into the genome of Flipin-TRex ™-293 cells (Invitrogen). Selection with Hygromycin and single clone isolation was followed by a test for Zeocin™-sensitivity to ensure proper recombination into a single locus in the genome. Cells were maintained in Doxycycline unless otherwise specified. YAPsh and control constructs were generously provided by Taran Gujral (Gujral and Kirschner, 2017). After transfection and antibiotic selection for two weeks, isolated single colonies were observed and were individually isolated. To label cells with nuclear localized mCherry, nuclear-mCherry was excised from pBRY-nuclear mCherry-IRES-PURO (Addgene plasmid #52409) by digestion with EcoRI and NotI and ligated into lentiviral expression plasmid pLVX-Ef1α-N1-mCherry. Lentiviruses were then produced via Fugene 6 co-transfection of psPAX2 (Addgene plasmid #12260) and pMD2.G (Addgene plasmid #12259) into 293T cells. One week after cells were infected with Lentivirus, they were FACS sorted for mCherry expression.

All cell lines were maintained in DMEM (Gibco, 10569010) supplemented with 10% FBS (Gibco), 1% P/S (Gibco). Neutralization antibodies against CYR61 (NOVUS biological) or CTGF (Peprotech) were used at the indicated concentrations in maintenance medium. Before growth factor treatment, cells were starved overnight in medium supplemented with 0.2% BSA instead of 10% FBS. Exogenous CYR61 and CTGF (Peprotech) were then added in medium with 10% FBS at 1ug/ml concentrations. All images were acquired from cells continuously cultured on 96-well plates (Eppendorf, unless otherwise specified) in an Incucyte Zoom apparatus installed in an incubator. Stiffness-dependent imaging was done on collagen-I coated 96-well plates (Matrigen). Transwell assays were done using 5mm Transwell^®^ plates with a 0.4 µm Pore Polycarbonate Membrane Insert (Corning). Recipient cells were cultured on the bottom of the plates while feeder cells were cultured on the insert.

### Automated image analysis

After image acquisition, red nuclei were segmented using Incucyte ZOOM’s automated image analysis software. All wells from the same plate were segmented using the same parameters as was all the data from the same time course. The resulting average area estimated for the objects in the red channel are what we refer to as the average nuclear area, while the number of the objects estimated per well were used as the number of nuclei.

### Total protein staining

Cells were trypsinized to a single-cell suspension, before they were thoroughly washed in PBS and fixed for 20 minutes in ice cold 4% PFA solution. Cells were then washed in PBS and permeabilized using 0.5% Triton-X for 10 minutes at room temperature. After repeated washes, cells were kept on ice with a 1:1000 dilution of Hoechst 3342 and 0.4 μg/ml of Alexa Fluor^®^ 647 Succinimidyl Ester (Invitrogen) in a light protected tube for 45 minutes before they were analyzed on an LSRII flow cytometer (BD Biosciences) at Harvard’s department of Systems Biology flow facility.

### RNA-Sequencing and data analysis

293 cells expressing Tet-inducible YAP5SA or nGFP were grown to confluence in medium supplemented with Tet-approved FBS (Takara) then supplemented with 0 or 1 ng/ml of Dox for the duration indicated. Total RNA was harvested and isolated using Qiagen’s RNeasy mini kit. Samples were further processed at Harvard’s Biopolymer facility as follows: After polyA enrichment of mRNA using Takara’s PrepX PolyA mRNA Isolation Kit, libraries were prepared using Illumina’s PrepX™ RNA-Seq Library reagent kit on the Apollo 324 system. Samples were pooled then single-end sequenced on the Illumina NextSeq 500 platform. Sequencing alignment was done using STAR (Dobin et al., 2013) and transcriptome assembly and differential expression was done using Cufflinks (Trapnell et al., 2012). Each comparison group contained at least two independent biological repeats. Data is deposited in GEO under accession number: xxx.

### Mass spectrometry and data analysis

Cells were kept under similar conditions as those used for RNA-sequencing. Cellular protein lysate was collected and snap frozen in urea-free lysis buffer supplemented with protease and phosphatase inhibitors (Roche). The supernatant was collected and filtered on a low protein-binding PVDF 0.22 μm membrane to eliminate cell debris, then snap frozen with protease inhibitors (Roche) for further processing. Lysates were reduced with 5 mM DTT, alkylated with 15 mM N-ethylmaleimide for 30 minutes in the dark, alkylation reactions quenched with 50 mM freshly prepared DTT and proteins precipitated by methanol/chloroform precipitation. Digests were carried out in 200 mM EPPS pH 8.5 in presence of 2% acetonitrile (v/v) with LysC (Wako, 2mg/ml, used 1:75) for 3 hours at room temperature and after subsequent addition of trypsin (Promega #V5111, stock 1:75) over night at 37°C.

Missed cleavage rate was assayed from a small aliquot by mass spectrometry. For whole proteome analysis, digests were directly labeled with TMT reagents (Thermo Fisher Scientific). Labeling efficiency and TMT ratios were assayed by mass spectrometry, while labeling reactions were stored at −80°C. After quenching of TMT labeling reactions with hydroxylamine, TMT labeling reactions were mixed, solvent evaporated to near completion and TMT labeled peptides purified and desalted by acidic reversed phase C18 chromatography. Peptides were then fractionated by alkaline reversed phase chromatography into 96 fractions and combined into 24 samples. Before mass spectrometric analysis, peptides were desalted over Stage Tips (Rappsilber et al., 2003).

Data were collected by a MultiNotch SPS MS^3^ method (McAlister et al., 2014) using an Orbitrap Lumos mass spectrometer (Thermo Fisher Scientific) coupled to a Proxeon EASY-nLC 1000 liquid chromatography (LC) system (Thermo Fisher Scientific). The 100 μm inner diameter capillary column used was packed with C18 resin (Accucore 2.6 μm, 150 Å, Thermo Fisher Scientific). Peptides of each fraction were separated over 3-5-hour acidic acetonitrile gradients by LC prior to mass spectrometry (MS) injection. The first sequence scan was an MS1 spectrum (Orbitrap analysis; resolution 120,000; mass range 400−1400 Th). MS2 analysis followed collision-induced dissociation (CID, CE=35) with a maximum ion injection time of 150 ms and an isolation window of 0.7 Da. To obtain quantitative information, MS^3^ precursors were fragmented by high-energy collision-induced dissociation (HCD) and analyzed in the Orbitrap at a resolution of 50,000 at 200 Th. MS^3^ injection time for phosphopeptides was 150 and 200 ms at a resolution of 50,000. Further details on LC and MS parameters and settings used were described recently (Paulo et al., 2016a).

Peptides were searched with a SEQUEST-based in-house software against a human database with a target decoy database strategy and a false discovery rate (FDR) of 1% set for peptide-spectrum matches following filtering by linear discriminant analysis (LDA) and a final collapsed protein-level FDR of 1%. Quantitative information on peptides was derived from MS^3^ scans. Quant tables were generated requiring an MS2 isolation specificity of >75% for each peptide and a sum of TMT s/n of >0 over all channels for any given peptide and exported as TAB-separated raw text data. The peptide-level sums of TMT s/n signal were integrated into protein-level and converted into relative ratio with 90% confidence intervals using a recently developed method BACIQ - a Bayesian approach to confidence inference for quantitative proteomics. Briefly, this approach reconciles disagreement across multiple peptides and differences in absolute levels of peptide signal, reporting the ratios across conditions for a given protein, modeling quantitative proteomics measurement as a random variable distributed according to hierarchical Dirichlet-Multinomial probability distribution. This method accounts for the absolute level of the peptide signal in a way calibrated for a particular MS instrument. More specifically we used a charge conversion value of 1.7 as previously fitted by us for this instrument and mass resolution (Peshkin et al., 2017). Details of the TMT intensity quantification method and further search parameters applied were described in (Paulo et al., 2016b).

Gene ontology analysis was done using DAVID (Huang et al., 2009) using all identified proteins as background.

## References

Anzi, S., Stolovich-Rain, M., Klochendler, A., Fridlich, O., Helman, A., Paz-Sonnenfeld, A., Avni-Magen, N., Kaufman, E., Ginzberg, M.B., Snider, D., et al. (2018). Postnatal Exocrine Pancreas Growth by Cellular Hypertrophy Correlates with a Shorter Lifespan in Mammals. Dev. Cell 45, 726–737.e3.

Basu, S., Totty, N.F., Irwin, M.S., Sudol, M., and Downward, J. (2003). Akt phosphorylates the Yes-associated protein, YAP, to induce interaction with 14-3-3 and attenuation of p73-mediated apoptosis. Mol. Cell 11, 11–23.

Chang, T.-Y., Chen, C., Lee, M., Chang, Y.-C., Lu, C.-H., Lu, S.-T., Wang, D.-Y., Wang, A., Guo, C.-L., and Cheng, P.-L. (2017). Paxillin facilitates timely neurite initiation on soft-substrate environments by interacting with the endocytic machinery. ELife 6.

Chen, C.C., Mo, F.E., and Lau, L.F. (2001). The angiogenic factor Cyr61 activates a genetic program for wound healing in human skin fibroblasts. J. Biol. Chem. 276, 47329–47337.

Diril, M.K., Ratnacaram, C.K., Padmakumar, V.C., Du, T., Wasser, M., Coppola, V., Tessarollo, L., and Kaldis, P. (2012). Cyclin-dependent kinase 1 (Cdk1) is essential for cell division and suppression of DNA re-replication but not for liver regeneration. Proc. Natl. Acad. Sci. U. S. A. 109, 3826–3831.

Dobin, A., Davis, C.A., Schlesinger, F., Drenkow, J., Zaleski, C., Jha, S., Batut, P., Chaisson, M., and Gingeras, T.R. (2013). STAR: ultrafast universal RNA-seq aligner. Bioinformatics 29, 15–21.

Dong, J., Feldmann, G., Huang, J., Wu, S., Zhang, N., Comerford, S.A., Gayyed, M.F., Anders, R.A., Maitra, A., and Pan, D. (2007). Elucidation of a universal size-control mechanism in Drosophila and mammals. Cell 130, 1120–1133.

Dupont, S., Morsut, L., Aragona, M., Enzo, E., Giulitti, S., Cordenonsi, M., Zanconato, F., Le Digabel, J., Forcato, M., Bicciato, S., et al. (2011). Role of YAP/TAZ in mechanotransduction. Nature 474, 179–183.

Elosegui-Artola, A., Andreu, I., Beedle, A.E.M., Lezamiz, A., Uroz, M., Kosmalska, A.J., Oria, R., Kechagia, J.Z., Rico-Lastres, P., Le Roux, A.-L., et al. (2017). Force Triggers YAP Nuclear Entry by Regulating Transport across Nuclear Pores. Cell 171, 1397–1410.e14.

Ernst, S., Demirci, C., Valle, S., Velazquez-Garcia, S., and Garcia-Ocaña, A. (2011). Mechanisms in the adaptation of maternal β-cells during pregnancy. Diabetes Manag. Lond. Engl. 1, 239–248.

Fankhauser, G. (1945). Maintenance of normal structure in heteroploid salamander larvae, through compensation of changes in cell size by adjustment of cell number and cell shape. J. Exp. Zool. 100, 445–455.

Gilgenkrantz, H., and Collin de l’Hortet, A. (2018). Understanding Liver Regeneration: From Mechanisms to Regenerative Medicine. Am. J. Pathol. 188, 1316–1327.

Ginzberg, M.B., Chang, N., D’Souza, H., Patel, N., Kafri, R., and Kirschner, M.W. (2018). Cell size sensing in animal cells coordinates anabolic growth rates and cell cycle progression to maintain cell size uniformity. ELife 7.

Groupe, V., and Manaker, R.A. (1956). Discrete foci of altered chicken embryo cells associated with Rous sarcoma virus in tissue culture. Virology 2, 838–840.

Gujral, T.S., and Kirschner, M.W. (2017). Hippo pathway mediates resistance to cytotoxic drugs. Proc. Natl. Acad. Sci. U. S. A. 114, E3729–E3738.

Halder, G., Dupont, S., and Piccolo, S. (2012). Transduction of mechanical and cytoskeletal cues by YAP and TAZ. Nat. Rev. Mol. Cell Biol. 13, 591–600.

Hansen, C.G., Ng, Y.L.D., Lam, W.-L.M., Plouffe, S.W., and Guan, K.-L. (2015). The Hippo pathway effectors YAP and TAZ promote cell growth by modulating amino acid signaling to mTORC1. Cell Res. 25, 1299–1313.

Holbourn, K.P., Acharya, K.R., and Perbal, B. (2008). The CCN family of proteins: structure–function relationships. Trends Biochem. Sci. 33, 461–473.

Huang, D.W., Sherman, B.T., and Lempicki, R.A. (2009). Systematic and integrative analysis of large gene lists using DAVID bioinformatics resources. Nat. Protoc. 4, 44–57.

Kamińska, K., Szczylik, C., Bielecka, Z.F., Bartnik, E., Porta, C., Lian, F., and Czarnecka, A.M. (2015). The role of the cell–cell interactions in cancer progression. J. Cell. Mol. Med. 19, 283–296.

Katsube, K., Sakamoto, K., Tamamura, Y., and Yamaguchi, A. (2009). Role of CCN, a vertebrate specific gene family, in development. Dev. Growth Differ. 51, 55–67.

Kim, N.-G., and Gumbiner, B.M. (2015). Adhesion to fibronectin regulates Hippo signaling via the FAK-Src-PI3K pathway. J. Cell Biol. 210, 503–515.

Lane, S.W., Williams, D.A., and Watt, F.M. (2014). Modulating the stem cell niche for tissue regeneration. Nat. Biotechnol. 32, 795–803.

Latinkic, B.V., Mercurio, S., Bennett, B., Hirst, E.M.A., Xu, Q., Lau, L.F., Mohun, T.J., and Smith, J.C. (2003). Xenopus Cyr61 regulates gastrulation movements and modulates Wnt signalling. Dev. Camb. Engl. 130, 2429–2441.

Leask, A., and Abraham, D.J. (2006). All in the CCN family: essential matricellular signaling modulators emerge from the bunker. J. Cell Sci. 119, 4803–4810.

Lee, M.-J., Byun, M.R., Furutani-Seiki, M., Hong, J.-H., and Jung, H.-S. (2014). YAP and TAZ regulate skin wound healing. J. Invest. Dermatol. 134, 518–525.

Lu, L., Finegold, M.J., and Johnson, R.L. (2018). Hippo pathway coactivators Yap and Taz are required to coordinate mammalian liver regeneration. Exp. Mol. Med. 50, e423.

McAlister, G.C., Nusinow, D.P., Jedrychowski, M.P., Wühr, M., Huttlin, E.L., Erickson, B.K., Rad, R., Haas, W., and Gygi, S.P. (2014). MultiNotch MS^3^ enables accurate, sensitive, and multiplexed detection of differential expression across cancer cell line proteomes. Anal. Chem. 86, 7150–7158.

Paulo, J.A., O’Connell, J.D., Everley, R.A., O’Brien, J., Gygi, M.A., and Gygi, S.P. (2016a). Quantitative mass spectrometry-based multiplexing compares the abundance of 5000 S. cerevisiae proteins across 10 carbon sources. J. Proteomics 148, 85–93.

Paulo, J.A., O’Connell, J.D., and Gygi, S.P. (2016b). A Triple Knockout (TKO) Proteomics Standard for Diagnosing Ion Interference in Isobaric Labeling Experiments. J. Am. Soc. Mass Spectrom. 27, 1620–1625.

Penzo-Méndez, A.I., and Stanger, B.Z. (2015). Organ-Size Regulation in Mammals. Cold Spring Harb. Perspect. Biol. 7, a019240.

Peshkin, L., Ryazanova, L., and Wuhr, M. (2017). Bayesian Confidence Intervals for Multiplexed Proteomics Integrate Ion-Statistics with Peptide Quantification Concordance. BioRxiv.

Pierre François Verhulst (1845). Recherches mathématiques sur la loi d’accroissement de la population. Nouv Mém Acad. R. Sci B.-Lett. Brux. 1–41.

Rappsilber, J., Ishihama, Y., and Mann, M. (2003). Stop and go extraction tips for matrix-assisted laser desorption/ionization, nanoelectrospray, and LC/MS sample pretreatment in proteomics. Anal. Chem. 75, 663–670.

Robert A. Weinberg (2007). The Biology of Cancer. (New York: Garland Science), pp. 61–65.

Shu, Z., and Deng, W.-M. (2017). Differential Regulation of Cyclin E by Yorkie-Scalloped Signaling in Organ Development. G3 Bethesda Md 7, 1049–1060.

Stoker, M.G., and Rubin, H. (1967). Density dependent inhibition of cell growth in culture. Nature 215, 171–172.

Tapon, N., Harvey, K.F., Bell, D.W., Wahrer, D.C.R., Schiripo, T.A., Haber, D.A., and Hariharan, I.K. (2002). salvador Promotes both cell cycle exit and apoptosis in Drosophila and is mutated in human cancer cell lines. Cell 110, 467–478.

Tian, X., Azpurua, J., Hine, C., Vaidya, A., Myakishev-Rempel, M., Ablaeva, J., Mao, Z., Nevo, E., Gorbunova, V., and Seluanov, A. (2013). High-molecular-mass hyaluronan mediates the cancer resistance of the naked mole rat. Nature 499, 346–349.

Trapnell, C., Roberts, A., Goff, L., Pertea, G., Kim, D., Kelley, D.R., Pimentel, H., Salzberg, S.L., Rinn, J.L., and Pachter, L. (2012). Differential gene and transcript expression analysis of RNA-seq experiments with TopHat and Cufflinks. Nat. Protoc. 7, 562–578.

Tsoularis, A., and Wallace, J. (2002). Analysis of logistic growth models. Math. Biosci. 179, 21–55.

Winick, M. (1968). Changes in nucleic acid and protein content of the human brain during growth. Pediatr. Res. 2, 352–355.

Xin, M., Kim, Y., Sutherland, L.B., Qi, X., McAnally, J., Schwartz, R.J., Richardson, J.A., Bassel-Duby, R., and Olson, E.N. (2011). Regulation of insulin-like growth factor signaling by Yap governs cardiomyocyte proliferation and embryonic heart size. Sci. Signal. 4, ra70.

Yimlamai, D., Christodoulou, C., Galli, G.G., Yanger, K., Pepe-Mooney, B., Gurung, B., Shrestha, K., Cahan, P., Stanger, B.Z., and Camargo, F.D. (2014). Hippo pathway activity influences liver cell fate. Cell 157, 1324–1338.

Zanconato, F., Forcato, M., Battilana, G., Azzolin, L., Quaranta, E., Bodega, B., Rosato, A., Bicciato, S., Cordenonsi, M., and Piccolo, S. (2015). Genome-wide association between YAP/TAZ/TEAD and AP-1 at enhancers drives oncogenic growth. Nat. Cell Biol. 17, 1218–1227.

Zhang, J., Ji, J.-Y., Yu, M., Overholtzer, M., Smolen, G.A., Wang, R., Brugge, J.S., Dyson, N.J., and Haber, D.A. (2009). YAP-dependent induction of amphiregulin identifies a non-cell-autonomous component of the Hippo pathway. Nat. Cell Biol. 11, 1444–1450.

Zhao, B., Wei, X., Li, W., Udan, R.S., Yang, Q., Kim, J., Xie, J., Ikenoue, T., Yu, J., Li, L., et al. (2007). Inactivation of YAP oncoprotein by the Hippo pathway is involved in cell contact inhibition and tissue growth control. Genes Dev. 21, 2747–2761.

Zhao, B., Li, L., Tumaneng, K., Wang, C.-Y., and Guan, K.-L. (2010). A coordinated phosphorylation by Lats and CK1 regulates YAP stability through SCF(beta-TRCP). Genes Dev. 24, 72–85.

