## Supplementary material for "YAP independently regulates cell size and population growth dynamics via non-cell autonomous mediators"

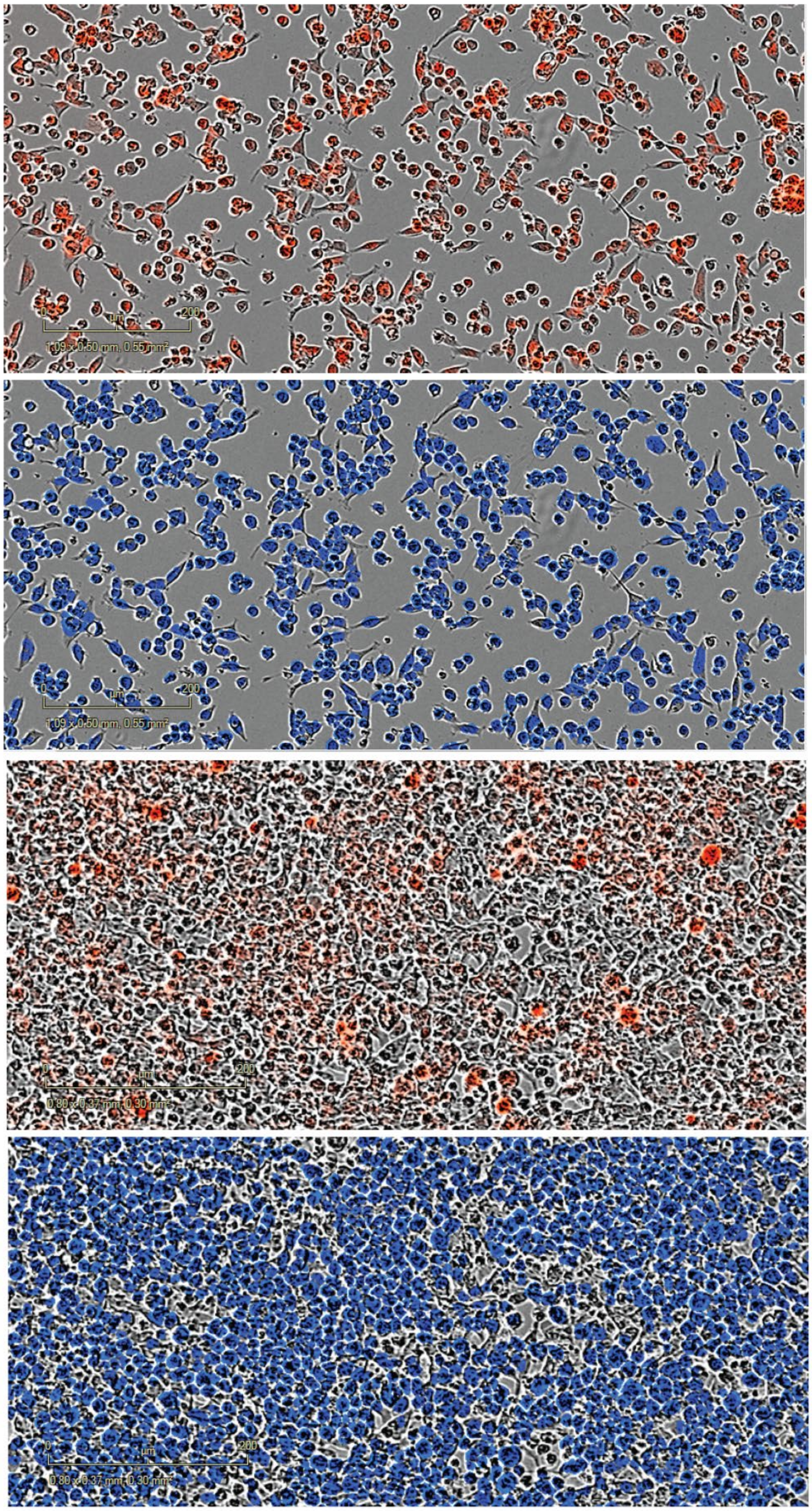


A — **50% confluence nmCherry signal**

B — **50% confluence nmCherry mask**

C — **100% confluence nmCherry signal**

D — **100% confluence nmCherry mask**

**Supplementary Figure S2: Example of images acquired on and analyzed by the Incucyte ZOOM to obtain data about the number of nuclei and their average area.** (A) HEK293 cells labelled with nmCherry growing at ~50% confluence. (B) The same field of view in A displaying the nmCherry mask used for estimating nuclear area and count as estimated by the Incucyte image analysis software. (C) Same as (A) at ~100% confluence. (D) Same as (B) at 100% confluence.

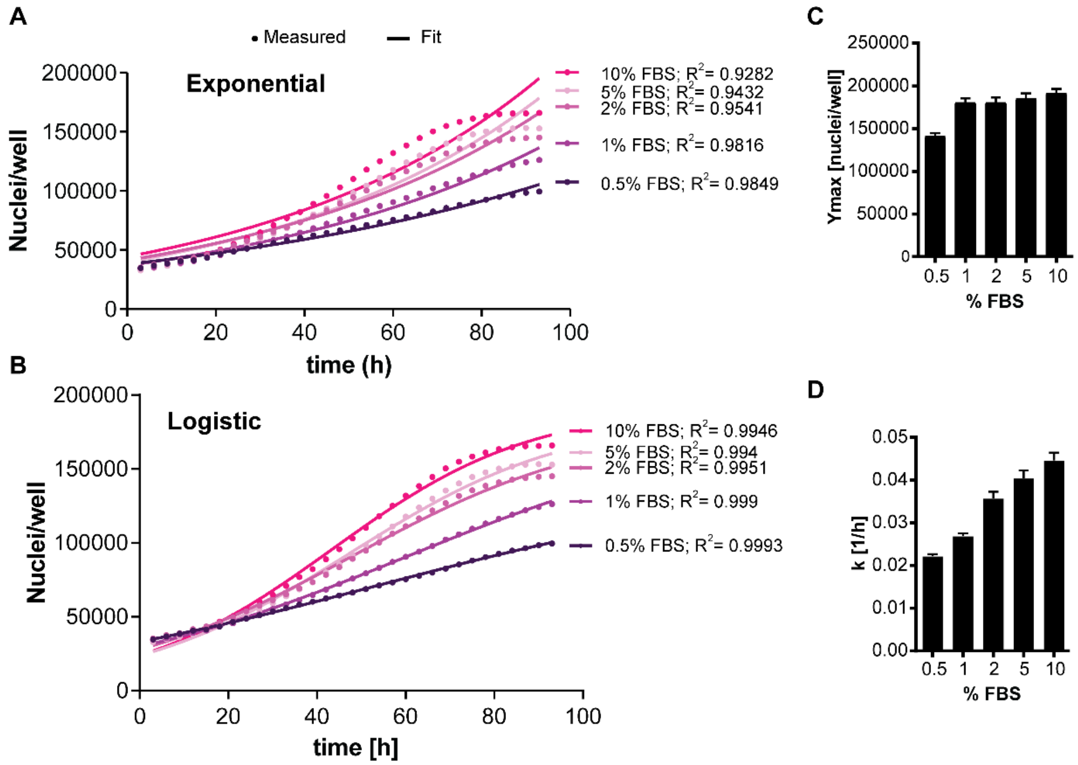


E

$$N(t) = \frac{Ymax * No}{No + \; e^{-kt}(Ymax - No)}$$

**Supplementary Figure S3: Varying serum concentration alters the growth rate of HEK293 cells but not their carrying capacity in culture.** The growth rate of HEK293 cells decreases over time, and is better modelled as logistic rather than exponential growth (A vs. B). The carrying capacity of a culture (Ymax) as estimated by the logistic growth equation does not increase by increasing bovine serum concentration (FBS) above 1 % (C) while the rate of growth (k) does (D). (E) Logistic growth equation. (n=5, mean±SEM)

A

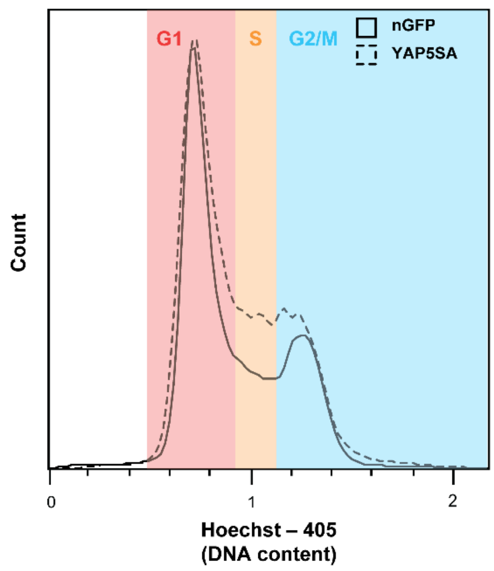


B

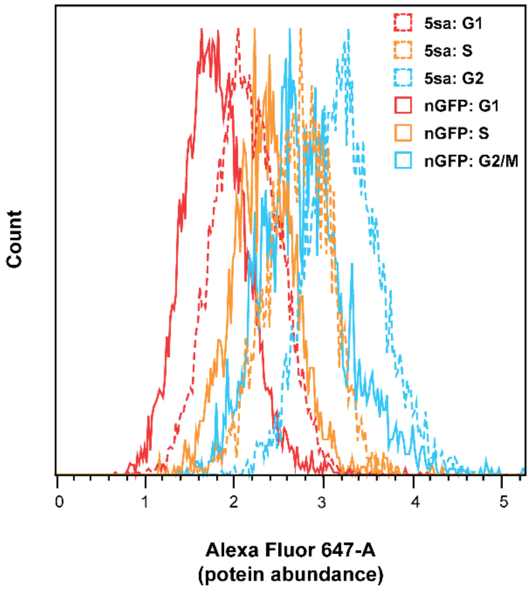


**Supplementary Figure S3: Cell cycle analysis and cell cycle dependent changes in protein content in YAP5SA vs nGFP controls.** (A) The fraction of cells in S and G2/M is higher in YAP5SA-expressing cells (YAP5SA) vs controls expressing nuclear GFP (nGFP). (B) Total protein content is higher in YAP5SA-expressing cells (5SA) vs nuclear GFP (nGFP) controls throughout the cell cycle.

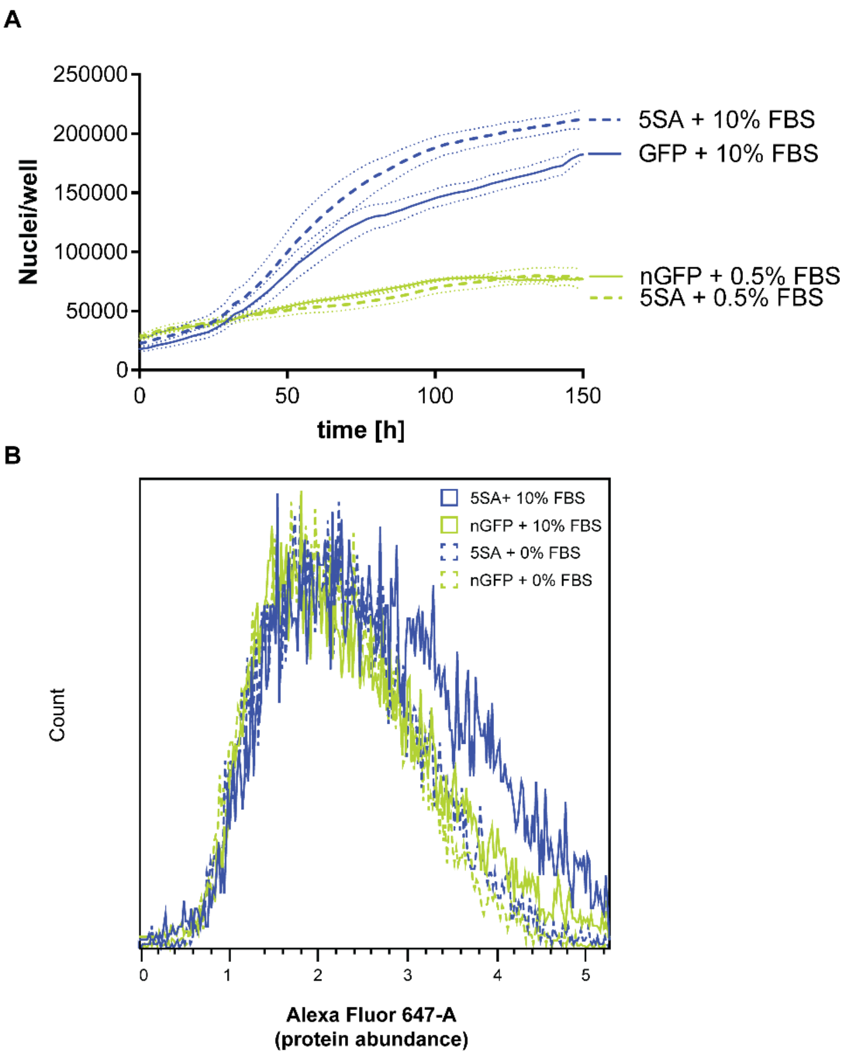


**Supplementary Figure S4: Serum is required for the 5SA-dependent changes in proliferation and size.** (A) Cell growth is reduced in the absence of serum, and is comparable between cells expressing nuclear GFP (nGFP) and YAP5SA (5SA) cells (n=5; mean±SEM). (B) Total protein content is higher in 5SA vs. nGFP controls in the presence of FBS, but not in its absence.

**Supplementary Table S5:** Table with protein and mRNA changes

**Supplementary Table S6:** Table with stiffness-dependent in gene expression after reanalysis of samples from GEO dataset GSE102350.
